# Fully Automated Peptide Mapping Protocol for Multi-Attribute Method by Liquid Chromatography–Tandem Mass Spectroscopy with a High-Throughput Robotic Liquid Handling System

**DOI:** 10.1101/2020.01.10.902338

**Authors:** Chen Qian, Ben Niu, Rod Brian Jimenez, Jihong Wang, Methal Albarghouthi, Xiaoyu Chen

## Abstract

The multi-attribute method (MAM) based on liquid chromatography–tandem mass spectroscopy is emerging as a powerful tool to directly monitor multiple product quality attributes simultaneously. Preparation of samples for MAM, however, is labor intensive, involving protein denaturation, disulfide bond reduction, free cysteine alkylation, and enzymatic digestion steps, which require significant analyst hands-on time while limiting result turnaround. Such complexity can also render nontrivial variations across analysts and laboratories. We describe the development of a fully automated peptide mapping procedure with a high-throughput robotic liquid handling system to improve sample handling capability and outcome reproducibility while saving analyst hands-on time. The automated procedure is completely hands-free, and setup requires the analyst only to prenormalize the sample concentrations and load buffers and reagents at their designated positions on the robotic deck. The robotic liquid handler performs all the subsequent preparation steps and stores the digested samples on a chiller unit to await retrieval. The convenience and flexibility provided by this automated peptide mapping method provides substantial benefits over manual sample preparation protocols. The optimized, automated procedure showed good reproducibility and results that were comparable to those of the manual procedure with respect to sequence coverage, digestion completeness, and quantification of posttranslational modifications. With this increased throughput, coupled with fast MAM analysis, more comprehensive characterization can be achieved.

## INTRODUCTION

With the emergence of an increasing number of biologic agents such as monoclonal antibodies (mAbs) and other engineered antibodies on the market, the demand for cost-effective, fast-turnaround process development and product characterization has driven technological advancements in the development of biopharmaceutical agents [1-6]. At the same time, regulatory agencies are promoting a “quality by design” (QbD) approach to the development process so that biologics will more closely adhere to targeted safety and efficacy profiles. In this approach, a quality target product profile (QTPP) is designed that encompasses the critical quality attributes (CQAs) of the biologic, and control strategies are developed and implemented to ensure that the QTPP is achieved in the end product [7-10]. During the manufacturing process, biologics are subjected to a number of posttranslational modifications (PTMs), including deamidation, oxidation, isomerization, and clipping [11-15]. Effective identification, monitoring, and control of these PTMs are critical to embrace the QbD approach [16-18].

Owing in part to the complexity of the quality attributes of biologics, the multi-attribute method (MAM) based on liquid chromatography (LC)–mass spectroscopy (MS) has debuted as an emerging technology, affording identification along with quantitative information on multiple product quality attributes throughout the development life cycle of biologics. It also aligns with QbD principles by making it possible to monitor the QTPP throughout the development process [19-23].

The typical protocol for preparing samples for MAM calls for denaturation, reduction, and alkylation, followed by tryptic digestion [24, 25, 20]. The generated tryptic peptides, which carry the product quality attributes, are then chromatographically separated and analyzed by MS. Major challenges to the use of MAM, however, lie in the variability of the sample preparation procedure and analytical instruments [26, 19], which can give rise to inconsistent results. To overcome instrument variability, instruments from the same vendor or the use of similar models is generally recommended [27]. A robust and consistent sample preparation procedure for MAM, however, is still lacking thus far. The currently used manual protocol of sample preparation is labor intensive and time consuming, severely limiting the throughput and turnaround of MAM. More important, due to the complexity of the manual procedure, variations across analysts, days, and laboratories can be significant, yielding inconsistent qualitative or quantitative information and creating challenges in the development of QTPPs and the implementation of control strategies. One solution is to automate the peptide mapping sample preparation procedure for MAM.

To the best of our knowledge, published reports on automation of peptide mapping procedures are scarce. Richardson et al. [25] reported on an automated, in-solution digestion protocol, using a commonly available high-performance LC (HPLC) autosampler. The protocol adapts a series of dilution steps, instead of desalting, to minimize the adverse effects of denaturants and reagents. Chelius et al. [28] reported an automated tryptic digestion procedure for a peptide mapping workflow, using Tecan with a custom-designed, 96-well desalting plate filled with silica-based size-exclusion (gel-filtration) medium. However, the need for customization of the 96-well desalting plate limited the applicability of their automated workflow. AssayMAP, recently developed by Agilent (Santa Clara, CA), is another automated platform used for analytical-scale protein sample preparation [29]. The peptide mapping procedure with AssayMAP, however, is not fully “walk-away” automation, as it requires manual intervention in the middle of the process, changing buffers on the deck before subsequent steps for buffer exchange and trypsin digestion.

Here we report the development and implementation of a fully walk-away automated procedure for peptide mapping, using the Hamilton STAR liquid handling platform with readily accessible labware, including the first-ever utilization of a 10K MWCO microdialysis plate in an automated process. Our results show that this automated procedure generated no artifact PTMs while affording high repeatability in terms of digestion efficiency and PTM quantitation. The increased throughput and reduced variation enabled by the automated workflow greatly expand the utilities of peptide mapping in process development and product characterization.

## MATERIALS AND METHODS

All biotherapeutic proteins, including the mAb, fragment crystallizable (Fc) fusion proteins and bispecifics, were expressed and purified at AstraZeneca (Gaithersburg, MD), using established protocols. All were stored at –80°C before analysis. Guanidine HCl (GdnHCl), dithiothreitol (DTT), iodoacetamide (IAM), hydrochloric acid, and the 10K MWCO microdialysis device were purchased from Thermo Fisher Scientific (Waltham, MA). PlusOne Urea was purchased from GE Healthcare Life Sciences (Chicago, IL). Tris(hydroxymethyl)aminomethane HCl solutions at different pHs were purchased from G-Biosciences (St. Louis, MO). Trypsin Gold MS Grade was purchased from Promega (Madison, WI).

Some of the automation consumables were sourced from Hamilton Company (Reno, NV) and included 50-µL and 300-µL conductive-filtered CO-RE tips and 50-mL reagent troughs. Starting sample material was contained in 2-mL, 96-square, deep-well plates sourced from Phenix Research Products (Candler, NC), and processing plates were round, U-bottomed microtiter plates with opaque lids, both sourced from Corning (Corning, NY). Eppendorf tubes of the 1.5-mL and 5-mL varieties were standard stock from VWR (Radnor, PA).

### Manual sample preparation for peptide mapping

Protein sample concentrations were diluted to 5.0 mg/mL with HPLC-grade water, followed by denaturation and reduction at 37°C for 30 min, using buffer comprising 7.2 M GdnHCl, 90 mM Tris, 0.1 mM ethylenediaminetetraacetic acid (EDTA), and 30 mM DTT, pH 7.4. Alkylation was performed by adding 500 mM IAM to a final concentration of 70 mM and incubating at room temperature for 30 minutes in the dark. This was followed by microdialysis to the buffer comprising 6 M urea and 150 mM Tris, pH 7.6, using a Pierce 96-well 10K MWCO microdialysis plate (catalog no. 88260; Thermo Fisher Scientific) according to the manufacturer’s instructions. Samples were then recovered from the microdialysis cassettes; concentrations were adjusted to 0.45 mg/mL with 100 mM Tris, pH 7.5; and Trypsin Gold (Promega) was added at a mass ratio of 1:12 (enzyme:sample) and incubated at 37°C. The digestion was quenched by adding 2% trifluoroacetic acid (TFA) (vol/vol) at room temperature.

### Automated sample preparation

Automated sample preparation was performed by using the STAR liquid handling robot (Hamilton Company). The robot is equipped with a 12-channel, independent spacing pipetting arm with a maximum transfer volume of 1 mL per channel, which was used for all the pipetting transfer steps. The STAR robot is also configured with a pair of CO-RE gripper paddles that leverage the independently spaced pipetting channels to move plates around the deck. The liquid handler deck included two peripherals: the Heater Shaker Unit (Hamilton Company) and an IC22 two-position heating/cooling block (Torrey Pines Scientific, Carlsbad, CA). Prior to robotic manipulation, the protein samples were normalized to a concentration of 5 mg/mL each with HPLC-grade water, and 50 µL of the resulting dilutions was aliquoted into the 96-well sample plate, which was loaded onto the deck (position A in Figure 1). The 96-well dialysis plate containing the dialysis cartridges submerged in dialysis buffer was placed at position D. All other required buffers and reagents were placed on the robotic deck as shown in Figure 1.

**Fig 1.**
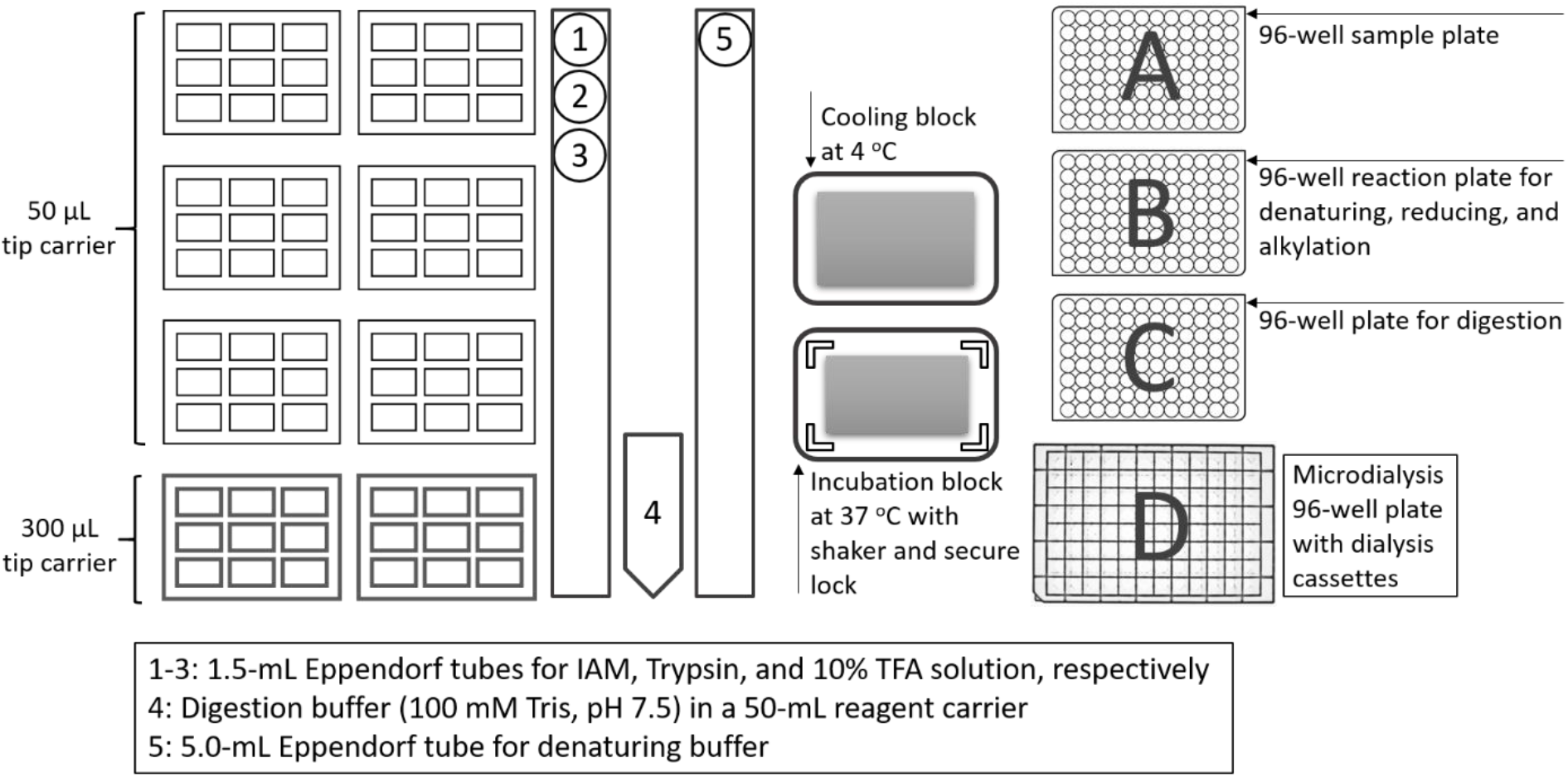
Schematic of the Hamilton Microlab STAR system for fully automated tryptic digestion procedure

A script written with Hamilton Venus Software mostly followed the manual sample preparation scheme, with a few slight variations to improve automatability. First, 44 µL of the denaturing buffer (7.2 M GdnHCl, 90 mM Tris, 0.1 mM EDTA, 30 mM DTT, pH 7.4) was transferred to an empty 96-well plate (position B in Figure 1). Next, 20 µL of each sample was transferred from the deep-well plate (position A in Figure 1) to the corresponding well of the denaturing buffer plate. The plate was then covered with an opaque black lid and moved to the heating block for 30 minutes of incubation at 37°C. After incubation, the plate was moved back to its original position and de-lidded, after which 10 µL of 500 mM IAM was added to each sample well. The denaturation plate was then re-lidded and subsequently incubated at room temperature for 30 minutes to alkylate free thiol groups. After alkylation, 35 µL of the alkylated sample was transferred to the dialysis plate, which was shaken at 500 rpm for 1 hour to desalt the samples. Toward the end of this shake time, 64 µL of digestion buffer and 6 µL of trypsin were pre-aliquoted into the other empty 96-well plate (position C in Figure 1), followed by the addition of dialyzed samples recovered after desalting was complete. The digestion buffer plate was then covered with another black opaque lid and moved to the heater for digestion at 37°C for 4 hours. Upon completion, the digestion plate was moved to the cooling block at 4°C, and 5 µL of 10% TFA was added to each sample well to quench the reaction. The plate containing the final quenched samples was covered with a lid and left on the cooler position, awaiting retrieval by an analyst.

### UPLC-MS/MS acquisition

For analysis by ultra-performance LC (UPLC)–tandem MS (MS/MS), samples were transferred to individual autosampler vials, or the whole plate was placed directly into a UPLC autosampler (Waters, Milford, MA). Aliquots (7 µg) of digested peptides were injected and separated on an Acquity UPLC Peptide BEH C18 Column (130 Å, 1.7 μm, 2.1 mm × 150 mm; Waters), using a linear gradient from 0% B to 40% B over 90 minutes. Mobile phase A contained 0.02% TFA (vol/vol) in water, and mobile phase B was 0.02% TFA (vol/vol) in acetonitrile. The column temperature was maintained at 55°C and the flow rate was kept at 0.2 mL/min. The digested samples were analyzed by coupling a Q-Exactive mass spectrometer (Thermo Fisher Scientific) with the UPLC system. The Q-Exactive mass spectrometer was equipped with an electrospray ionization source, and spray voltage was set at 3.5 kV. The capillary temperature was 320°C, and the mass range was set at 250–2,000 m/z. Data were acquired in data-dependent acquisition (DDA) mode, and the fragmentation mass spectra were obtained by HCD.

### Data analysis

The acquired *.raw MS files were searched directly with Byonic software (Protein Metrics) against a custom-built database containing the sequences of the protein of interest. Search parameters were set to include tryptic cleavage sites (C-terminal of Arg, Lys) with any number of missed cleavages. The search tolerance window was 15 ppm for all precursor ions and 50 ppm for product ions. Maximum precursor mass was set to 15,000 Da. For PTMs, the alkylation of Cys-containing peptides with IAM, which adds a carbamidomethyl group, was included as a fixed modification in the search. All other PTMs were considered variable. Details of PTMs and glycopeptide considerations are described in the Supplemental Information.

Quantitation of PTMs was based on ion abundances represented on integrated extracted-ion chromatograms (XICs), which were analyzed in Byologic software (Protein Metrics). Manual verification was performed for XIC integration consistency.

## RESULTS AND DISCUSSION

### Optimization of microdialysis condition

The tryptic digestion of antibodies typically involves denaturation, reduction, alkylation, and clean-up steps prior to digestion. A high concentration of chaotrope (e.g., GdnHCl or urea) is typically used for effective denaturation [30, 31]. The trypsin activity, however, can be adversely affected by even a small amount of denaturant (e.g., <0.3 M GdnHCl) [32]), resulting in poor digestion efficiency and decreased profile reproducibility. Sample clean-up (e.g., by buffer exchange) immediately prior to tryptic digestion is crucial to eliminate unwanted chemicals and maintain a consistent, digestion-friendly solution environment. Alternatively, serial dilution steps can be performed so that the chaotrope concentration can be diluted to a level that does not inhibit trypsin activity. We evaluated several commonly used, commercially available sample clean-up devices, including NAP-5 columns, Amicon centrifugal filters (Sigma-Aldrich, St. Louis, MO), Slide-A-Lyzer dialysis cassettes (Thermo Fisher Scientific), and the 96-well 10K MWCO microdialysis plate, and compared the outcomes with those of the serial-dilution approach.

For quantitative evaluation of tryptic digestion efficiency, we focused on the relative abundance of tryptic peptides with 0 missed cleavage and calculated the digestion completeness percentage under each condition as:

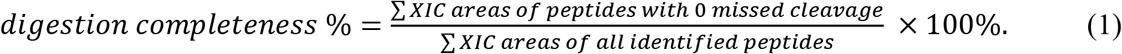

Both unmodified and modified peptides were considered. For peptides exhibiting multiple charge states, the XIC areas corresponded to the summation of integrals of each individual charge state.

Figure 2a shows the comparisons of digestion completeness percentage based on manual 3.5-hour tryptic digestion of NISTmAb with different cleanup approaches. Digestion completeness percentage reached approximately 90% when buffer exchange was performed, and the majority of product peptides had zero or one missed cleavage. However, digestion efficiency was significantly lower without buffer exchange (Figure 2a, “Serial dilution”), which resulted in approximately 25% of product peptides with two or more missed cleavages when diluted to a final concentration of 0.26 M GdnHCl. Indeed, we found numerous peptides with prominent missed cleavages in the Fc region, including the Fc glycopeptide with three missed cleavages (sequence TKPREEQYNSTYRVVSVLTVLHQDWLNGKEYK). In addition to digestion completeness percentage, we compared the sample recovery of each cleanup method as measured by UV-visible spectrophotometry of protein concentrations after cleanup. Both membrane-based approaches, i.e., using the microdialysis plate and Slide-A-Lyzer dialysis cassettes, resulted in noticeably higher sample recovery. After considering all of these factors, we adopted the 96-well microdialysis plate in the automated procedure because of its high sample capacity and readiness for automation workflow.

**Fig 2.**
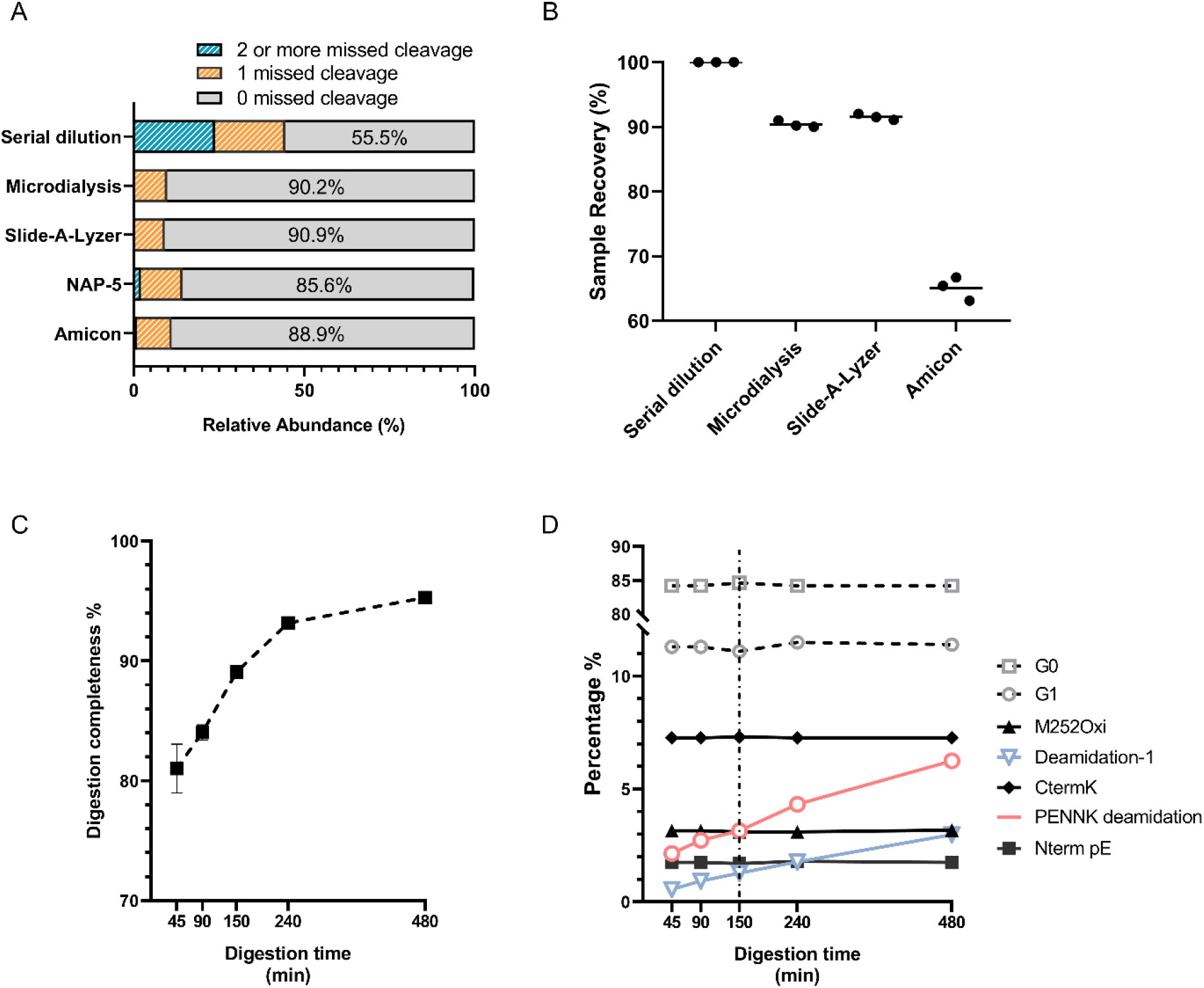
**a** Evaluation of trypsin digestion completeness using NISTmAb with different sample cleanup approaches prior to digestion. Without buffer exchange, the digestion completeness was compromised, as shown by the significantly increased extent of miscleavages in the serial dilution approach. **b** Evaluation of sample recovery using NISTmAb as measured by UV-visible spectrophotometry of protein concentrations after cleanup. Because the serial dilution approach was assumed to cause no sample loss, the recovery was set to 100%. **c** Tryptic digestion completeness percentage of mAb-A as a function of digestion time. Time points were 45, 90, 150, 240, and 480 minutes. **d** Monitoring of selected PQAs of mAb-A under each digestion time. (CtermK, C-terminal lysine; Nterm pE, N-terminal pyroglutamate)

To optimize the microdialysis conditions, four mAbs were manually tested under various microdialysis conditions. The parameters considered included dialysis duration and temperature, sample volume inside the cassette, and vortex speed. A screening Design of Experiments (DoE) was conducted first, using the NISTmAb to identify the factors that significantly influenced digestion completeness (see Supplementary Information for details). According to the DoE results, the dialysis duration and volume and the vortex speed had the largest absolute effect, whereas the effect of dialysis temperature was not significant (Supplementary Figure S1). The vortex speed during dialysis was positively correlated with digestion completeness, most likely due to improved buffer exchange efficiency with increased agitation. The sample volume of 35 μL inside the cassette also led to more efficient buffer exchange than a volume of 70 µL. Although longer microdialysis duration dramatically improved buffer exchange, the buffer exchange efficiency seemed to reach a plateau after 1 hour of dialysis.

### Optimization of digestion time

With the optimized microdialysis condition, we further assessed digestion completeness percentage as a function of digestion time. The time-dependent tryptic digestion samples were manually prepared at five different incubation times (45, 90, 150, 240, and 480 minutes) with four replicates at each time point. Our results (Figure 2c) suggest that, in the first 4 hours of digestion of mAb-A, the digestion completeness percentage was proportional to the digestion time, although the variation seemed to be greater at 45 minutes of digestion than at longer times. The increment of digestion completeness percentage, however, slowed down after 4 hours, probably because most peptides had been nearly fully digested after that time.

In parallel with the assessment of digestion completeness, the quantitation of PTMs at each time point was evaluated and reported (Figure 2d). Selected product quality attributes (PQAs) of mAb-A, including oxidation, deamidation, N-terminal cyclization, C-terminal Lys, and N-glycans, were monitored. Although the percentage of most PQAs remained unchanged regardless of the digestion time, we found two deamidations (one PENNYK peptide deamidation and one deamidation in the complementarity-determining region [CDR]) whose quantitation increased as a function of digestion time. The deamidation percentage of the PENNYK peptide, for example, reached 6.2% at 480 minutes of digestion and showed an almost linear increment of 0.5% per hour during the full span of digestion. This suggests that longer digestion time, despite allowing more thorough dissociation of peptides, can induce artifact modifications, and deamidation appears to be most susceptible, as the digestion is typically performed under mildly basic pH [14, 33]. We confirmed this artifact deamidation with low-pH tryptic digestion, in which the PENNYK deamidation percentage remained within 3–4% even after 24 hours of digestion [34]. To balance the digestion completeness and level of artifact modifications, in the automated procedure we adopted the digestion time of 150 minutes because of the reasonably good digestion completeness and minor artifact modifications that occurred with this time.

### Repeatability of automated procedure

The automated procedure closely resembled the manual procedure in its optimized microdialysis and digestion parameters. The most challenging aspect of mirroring the automated protocol to the manual procedure lay in the microdialysis procedure, given the small size of the opening on each individual dialysis cassette [35]. Therefore, calibration of automated movements for the pipetting steps during microdialysis was required to avoid mispositioning and ensure that sample recovery after microdialysis was comparable between the automated protocol and the manual protocol. (Calibration details are provided in the Supplemental Information.) Great consistency and comparability of sample recovery were achieved with a volume variability of less than 1 μL, as shown in Supplemental Figure S2.

To establish an automated protocol, however, several adjustments needed to be implemented. For example, to ensure the accuracy and reproducibility of liquid handling, the automated procedure was adapted to ensure that the minimal pipetting volume was no less than 5 μL. Because of this requirement, concentrations of stock solutions such as trypsin and TFA needed to be adjusted accordingly. To this end, to quench digestion, we added 10% TFA at a 1:20 volume ratio instead of pure TFA at a 1:50 volume ratio, given the reduced pipetting variability and the decreased volatility of less-concentrated TFA, which permits long-term TFA storage at room temperature. In addition, to guarantee effective mixing of solution during pipetting and dispensing, a mechanical mixing step was performed by repeatedly pipetting up and down after each dilution or reaction. Despite implementation of these adjustments in the automated protocol, we achieved largely consistent and comparable digestion profiles between the manual and automated procedures (Supplemental Figure S3).

This automated procedure was capable of handling a full 96-well plate of samples. A full plate run comprising different molecule modalities (including IgG1, IgG2, IgG4, Fc fusion protein, and bispecifics) was conducted. Figure 3a shows an overlay of UV chromatograms of six tryptic digestions of mAb-B (an IgG4 antibody) and the corresponding blank digestion with automation. The sequence coverage reached 99.7% for the heavy chain and 100% for the light chain. The digestion efficiency was high, as seen by the major species referring to peptides with 0 miscleavage, and the reproducibility of digestion was demonstrated by the consistent UV profile among the six samples, in which no new or missing peaks were observed. The maximal retention time difference was 0.1 minute for all peaks, with signals of more than 0.5% of the species with the highest UV signal (H14). A UV overlay of two other molecules (in six replicates), IgG1 and IgG2, are shown in Supplemental Figure S4. All peaks were aligned with comparable abundance across all the replicates.

**Fig 3.**
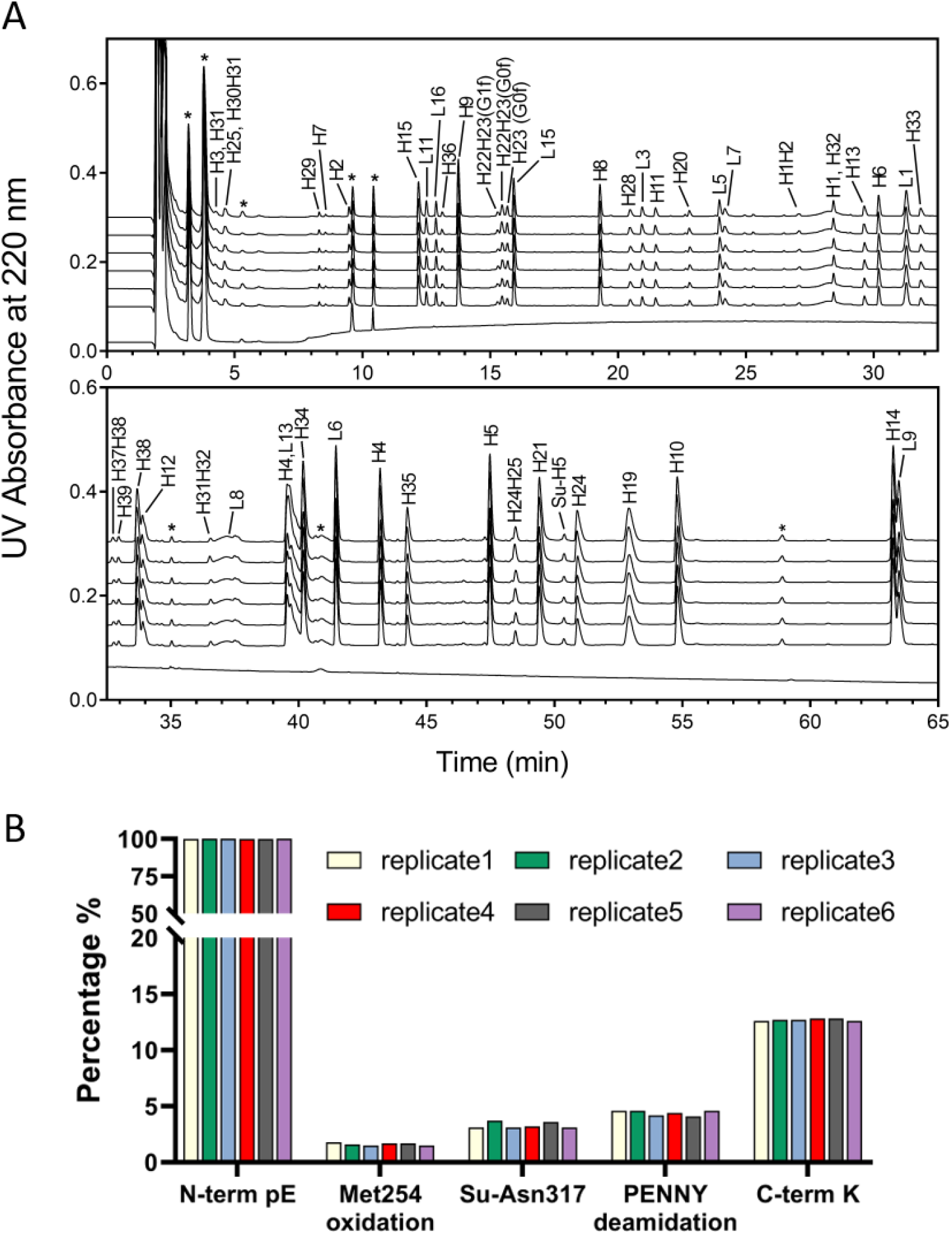
**a** UV chromatogram overlay of six tryptic digestions of mAb-B (an IgG4 antibody) and a blank digestion. Annotations indicate the heavy chain (H) and light chain (L) with the corresponding tryptic peptide numbers. Peaks labeled with asterisks (*) are artifact peaks from experimental reagents or trypsin autolysis. **b** PTM quantitation of selected attributes showed consistent levels of PTMs for the six tryptic digested mAb-B samples. C-term K, C-terminal lysine); (N-term pE, N-terminal pyroglutamate)

The PTM levels of the six mAb-B samples were also assessed (Figure 3b). The N terminal of the mAb-B heavy chain was fully processed, as seen by the approximately 100% pyroglutamate formation, whereas the C-terminal lysine of the mAb-B heavy chain was partially processed. The results showed good consistency of PTM quantitation across all six samples, with levels ranging from 1% to 100%; the maximum coefficient of variation was 8.4%.

To evaluate day-to-day variations in PTM quantitation, samples were prepared with the automated procedure in six replicates on three different days of three different weeks. Three in-house reference standard samples, in addition to the NISTmAb, were tested. For the NISTmAb, a set of PQAs, including N-terminal cyclization, Met255 oxidation, Asn364 deamidation, PENNY deamidation, C-terminal Lys, and G0F, were considered in the PTM quantitation. The results showed good repeatability for all monitored PQAs within each preparation and across different days, with a coefficient of variation of less than 10% (Figure 4a). Notably, these PTM quantitation results were also comparable to those obtained from manual sample preparation (data not shown).

**Fig 4.**
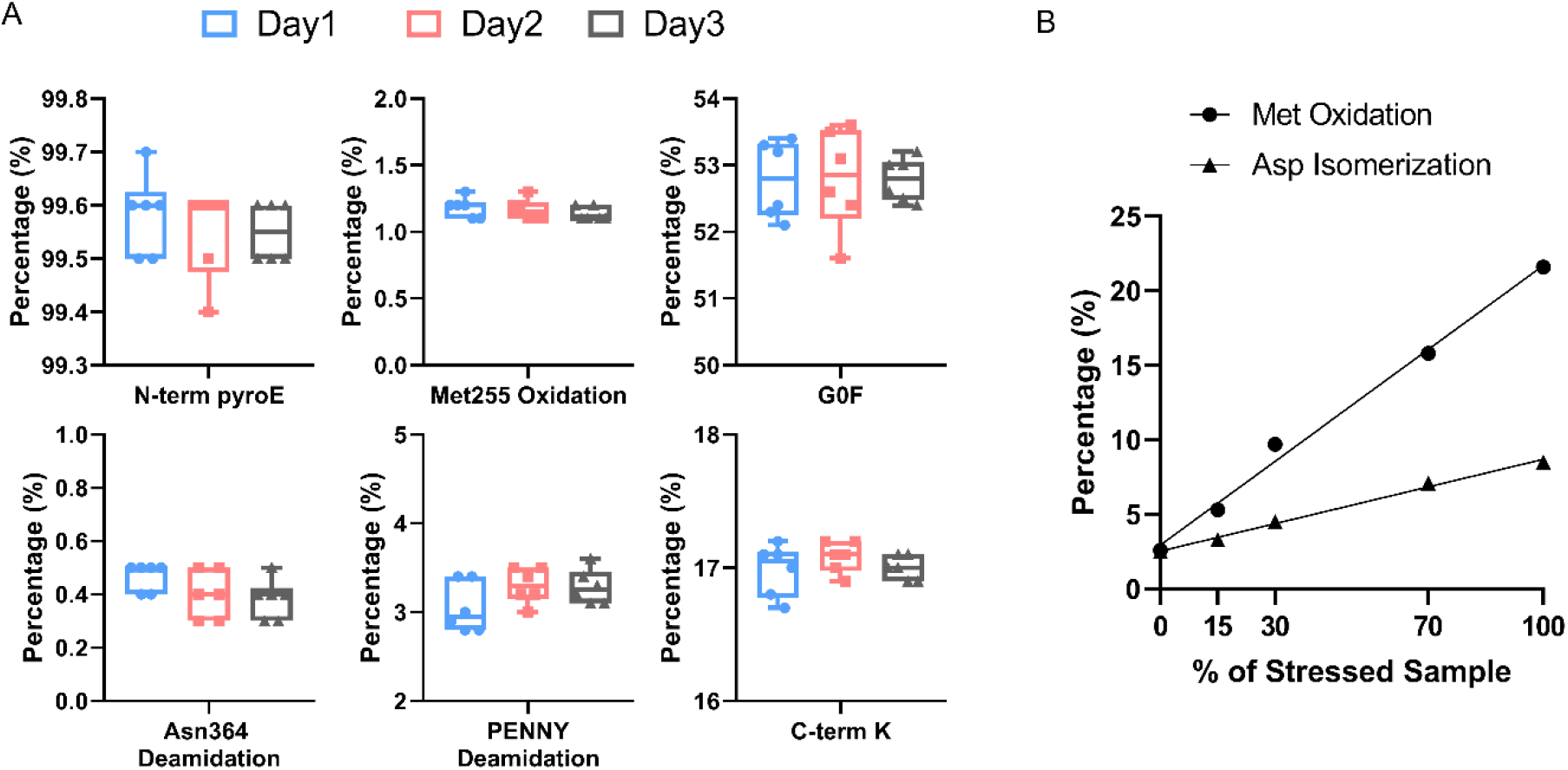
**a** Quantitation of selected PTMs of the NISTmAb demonstrated good repeatability of quantitation and minor day-to-day variations. **b** Stability-indicating PTMs of blended mAb-A samples measured as a function of blending ratio (0%, 15%, 30%, 70%, 100%) to demonstrate the quantitation linearity

We also evaluated the stability-indicating property of the automated preparation procedure. Nonstressed mAb-A drug substance samples were blended with degraded mAb-A drug product (60 months at 2–8°C storage) at five ratios: 0%, 15%, 30%, 70%, and 100% of the degraded drug product sample. The resulting blended samples were subjected to the automated preparation procedure and tested. The PTM percentage of Met252 oxidation and CDR isomerization were plotted as a function of the degraded drug product percentage (Figure 4b) to demonstrate the capability of linear quantitation for stability-indicating properties.

### Implementation for product CQA monitoring

The automated procedure significantly reduced the sample preparation time for analysts, and more importantly, it improved consistency across analysts, laboratories, and days, as the materials are fully commercially available and the operation settings are readily exportable. We have been using the automated procedure extensively for monitoring product quality attributes. Figure 5a shows the monitoring of the oxidation level of Met252 in a forced-oxidation study. With 50 ppm of H_2_O_2_ at 4°C, Met252 reached 49% oxidation after 243 hours of incubation. The quantitation of Met252 oxidation by this method was found to be comparable to that obtained with other orthogonal methods, including use of the QDa and ProteinA affinity column. The determined oxidation levels correlated well with the decrease of neonatal Fc receptor (FcRn) binding (Figure 5b), which is known to shorten serum half-life [12]. We further show that this automated procedure is applicable to process development, in which changes of mAb-A quality attributes during purification process can be easily monitored. A noticeable decrease in deamidation levels was observed for both CDR and Fc regions after the low-pH process step, whereas the Met252 level remained unchanged throughout all processes (Figure 5c).

**Fig 5.**
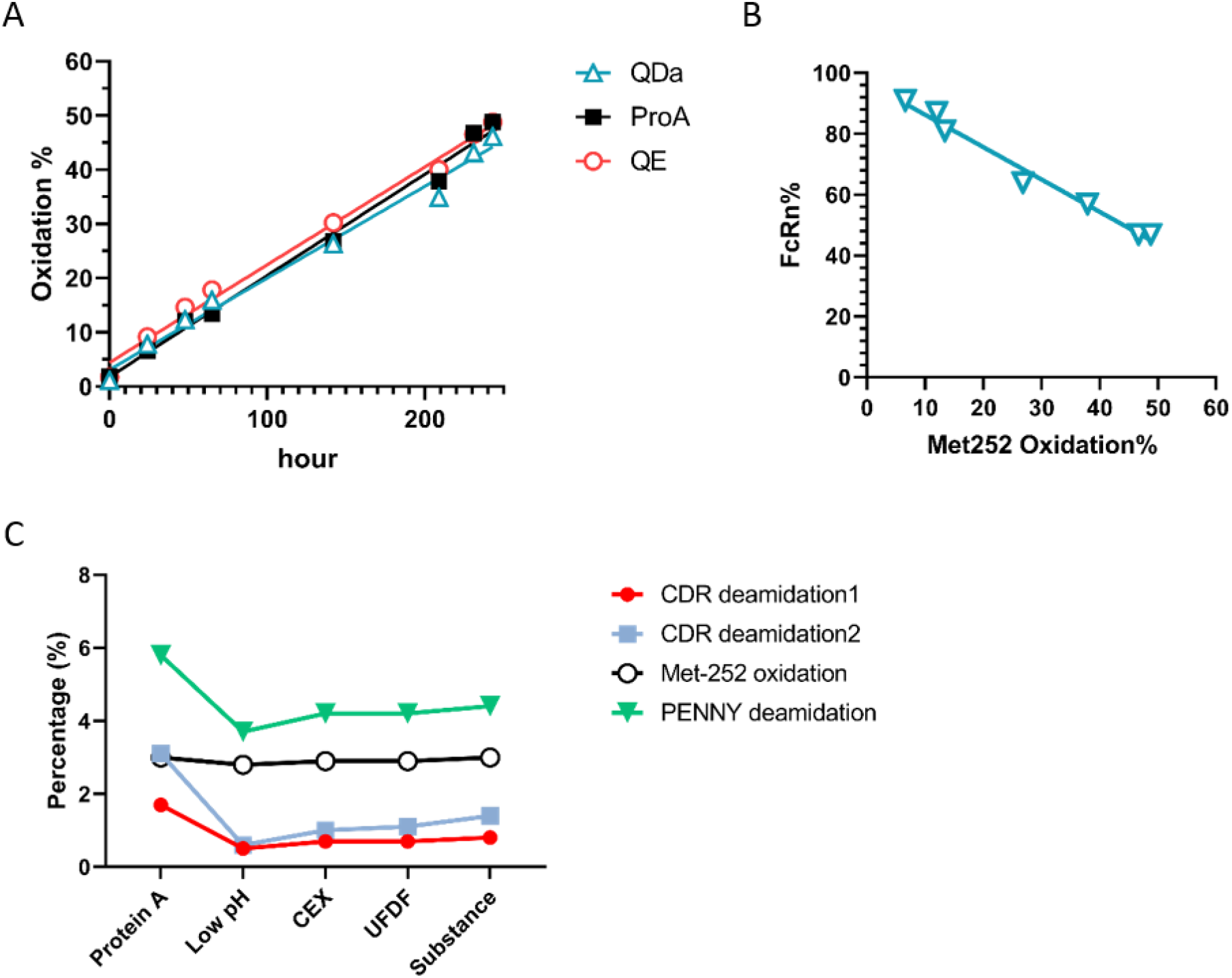
**a** Met252 oxidation levels measured by automated MAM in forced-oxidation study. The quantitation by automated MAM aligned well with results independently generated in QDa and ProteinA methods. **b** Correlation of the extent of Met252 oxidation and FcRn binding efficacy showed a linear relationship. **c** Use of automated MAM for monitoring several quality attributes in downstream purification process

## Conclusion

We report the development and optimization of a fully automated tryptic peptide mapping procedure on the Hamilton Star liquid handling system with buffer exchange by incorporating the commercially available 96-well microdialysis plate. This method demonstrated high digestion efficiency with good reproducibility, as indicated by consistency of both UV profiles and PTM quantitation. By implementing the automated protocol, the hands-on time of peptide mapping sample preparation was significantly reduced. The automated protocol demonstrated much smaller variations across analysts, laboratories, and days, and the implementation barrier was low because no complex training or domain-specific expertise was required to operate the system. Moreover, this automated procedure closely resembles the manual procedure in terms of reagent concentrations, pipetting volumes, and buffer exchange steps, affording UV chromatograms and interchangeable PTM quantitation that were comparable to those of the manual procedure. This automated protocol is currently routinely used to support product characterization and process development work. Future improvements of the procedure may include an automatic pre-adjustment of sample concentrations to accommodate samples that are provided in various concentrations.

## Acknowledgments

We thank Thomas Albanetti for the Hamilton basic training.

## Funding information

This study was supported by AstraZeneca.

**Compliance with ethical standards**

## REFERENCES

1. Buckley K, Ryder AG. Applications of raman spectroscopy in biopharmaceutical manufacturing: a short review. Appl Spectrosc. 2017;71:1085–116.

2. Cao M, De Mel N, Shannon A, Prophet M, Wang C, Xu W, Niu B, Kim J, Albarghouthi M, Liu D, Meinke E, Lin S, Wang X, Wang J. Charge variants characterization and release assay development for co-formulated antibodies as a combination therapy. mAbs. 2019;11:489–99.

3. Goswami D, Zhang J, Bondarenko PV, Zhang Z. MS-based conformation analysis of recombinant proteins in design, optimization and development of biopharmaceuticals. Methods. 2018;144:134–51.

4. Gronemeyer P, Ditz R, Strube J. Trends in upstream and downstream process development for antibody manufacturing. Bioengineering (Basel). 2014;1:188–212.

5. Kroll P, Hofer A, Ulonska S, Kager J, Herwig C. Model-based methods in the biopharmaceutical process lifecycle. Pharm Res. 2017;34:2596–613.

6. Valliere-Douglass J, Wallace A, Balland A. Separation of populations of antibody variants by fine tuning of hydrophobic-interaction chromatography operating conditions. J Chromatogr A. 2008;1214:81–9.

7. International Council for Harmonisation (2012) Development and Manufacture of Drug Substances (Q11). https://www.ich.org/products/guidelines/quality/article/quality-guidelines.html.

8. Kepert JF, Cromwell M, Engler N, Finkler C, Gellermann G, Gennaro L, Harris R, Iverson R, Kelley B, Krummen L, McKnight N, Motchnik P, Schnaible V, Taticek R. Establishing a control system using QbD principles. Biologicals. 2016;44:319–31.

9. Luciani F, Galluzzo S, Gaggioli A, Kruse NA, Venneugues P, Schneider CK, Pini C, Melchiorri D. Implementing quality by design for biotech products: are regulators on track? mAbs. 2015;7:451–5.

10. Yu LX, Amidon G, Khan MA, Hoag SW, Polli J, Raju GK, Woodcock J. Understanding pharmaceutical quality by design. AAPS J. 2014;16:771–83.

11. Blessy M, Patel RD, Prajapati PN, Agrawal YK. Development of forced degradation and stability indicating studies of drugs-A review. J Pharm Anal. 2014;4:159–65.

12. Gao X, Ji JA, Veeravalli K, Wang YJ, Zhang T, McGreevy W, Zheng K, Kelley RF, Laird MW, Liu J, Cromwell M. Effect of individual Fc methionine oxidation on FcRn binding: Met252 oxidation impairs FcRn binding more profoundly than Met428 oxidation. J Pharm Sci. 2015;104:368–77.

13. Harris RJ, Kabakoff B, Macchi FD, Shen FJ, Kwong M, Andya JD, Shire SJ, Bjork N, Totpal K, Chen AB. Identification of multiple sources of charge heterogeneity in a recombinant antibody. J Chromatogr B Biomed Sci Appl. 2001;752:233–45.

14. Pace AL, Wong RL, Zhang YT, Kao YH, Wang YJ. Asparagine deamidation dependence on buffer type, pH, and temperature. J Pharm Sci. 2013;102:1712–23.

15. Zhang YT, Hu J, Pace AL, Wong R, Wang YJ, Kao YH. Characterization of asparagine 330 deamidation in an Fc-fragment of IgG1 using cation exchange chromatography and peptide mapping. J Chromatogr B Analyt Technol Biomed Life Sci. 2014;965:65–71.

16. Beg S, Swain S, Rahman M, Hasnain MS, Imam SS (2019) Application of design of experiments in pharmaceutical product and process optimization. In: Beg S, Hasnain MS (eds) Pharmaceutical quality by design. Academic Press, New York, pp 43–64. doi:https://doi.org/10.1016/B978-0-12-815799-2.00003-4

17. Finkler C, Krummen L. Introduction to the application of QbD principles for the development of monoclonal antibodies. Biologicals. 2016;44:282–90.

18. Kona R, Fahmy RM, Claycamp G, Polli JE, Martinez M, Hoag SW. Quality-by-design III: application of near-infrared spectroscopy to monitor roller compaction in-process and product quality attributes of immediate release tablets. AAPS PharmSciTech. 2015;16:202–16.

19. Rogers RS, Abernathy M, Richardson DD, Rouse JC, Sperry JB, Swann P, Wypych J, Yu C, Zang L, Deshpande R. A view on the importance of “multi-attribute method” for measuring purity of biopharmaceuticals and improving overall control strategy. AAPS J. 2017;20:7.

20. Rogers RS, Nightlinger NS, Livingston B, Campbell P, Bailey R, Balland A. Development of a quantitative mass spectrometry multi-attribute method for characterization, quality control testing and disposition of biologics. mAbs. 2015;7:881–90.

21. Xu W, Jimenez RB, Mowery R, Luo H, Cao M, Agarwal N, Ramos I, Wang X, Wang J. A quadrupole Dalton-based multi-attribute method for product characterization, process development, and quality control of therapeutic proteins. mAbs. 2017;9:1186–96.

22. Xu XQ, H; Li, N. LC-MS multi-attribute method for characterization of biologics. J Appl Bioanal. 2017;3:21–5.

23. Zhang Y, Guo J. Characterization and QC of biopharmaceuticals by MS-based ‘multi-attribute method’: advantages and challenges. Bioanalysis. 2017;9:499–502.

24. Mouchahoir T, Schiel JE. Development of an LC-MS/MS peptide mapping protocol for the NISTmAb. Anal Bioanal Chem. 2018;410:2111–26.

25. Richardson J, Shah B, Xiao G, Bondarenko PV, Zhang Z. Automated in-solution protein digestion using a commonly available high-performance liquid chromatography autosampler. Anal Biochem. 2011;411:284–91.

26. Hada V, Bagdi A, Bihari Z, Timari SB, Fizil A, Szantay C, Jr. Recent advancements, challenges, and practical considerations in the mass spectrometry-based analytics of protein biotherapeutics: a viewpoint from the biosimilar industry. J Pharm Biomed Anal. 2018;161:214–38.

27. Zhang Z, Shah B, Guan X. Reliable LC-MS multiattribute method for biotherapeutics by run-time response calibration. Anal Chem. 2019;91:5252–60.

28. Chelius D, Xiao G, Nichols AC, Vizel A, He B, Dillon TM, Rehder DS, Pipes GD, Kraft E, Oroska A, Treuheit MJ, Bondarenko PV. Automated tryptic digestion procedure for HPLC/MS/MS peptide mapping of immunoglobulin gamma antibodies in pharmaceutics. J Pharm Biomed Anal. 2008;47:285–94.

29. Beban M, Tang N, van den Heuvel Z, Knorr D. Antibody purification with AssayMAP, a high-throughput microchromatography platform. Journal of Biomolecular Techniques. 2012;23:S27–S8.

30. Ren D, Pipes GD, Liu D, Shih LY, Nichols AC, Treuheit MJ, Brems DN, Bondarenko PV. An improved trypsin digestion method minimizes digestion-induced modifications on proteins. Anal Biochem. 2009;392:12–21.

31. Salvi G, De Los Rios P, Vendruscolo M. Effective interactions between chaotropic agents and proteins. Proteins. 2005;61:492–9.

32. Proc JL, Kuzyk MA, Hardie DB, Yang J, Smith DS, Jackson AM, Parker CE, Borchers CH. A quantitative study of the effects of chaotropic agents, surfactants, and solvents o the digestion efficiency of human plasma proteins by trypsin. J Proteome Res. 2010;9:5422–37.

33. Peters B, Trout BL. Asparagine deamidation: pH-dependent mechanism from density functional theory. Biochemistry. 2006;45:5384–92.

34. Cao M, Xu W, Niu B, Kabundi I, Luo H, Prophet M, Chen W, Liu D, Saveliev SV, Urh M, Wang J. An automated and qualified platform method for site-specific succinimide and deamidation quantitation using low-pH peptide mapping. J Pharm Sci. 2019;108:3540–9.

35. Thermo Scientific (2019) Pierce 96-well Microdialysis Plate, 10K MWCO. https://www.thermofisher.com/order/catalog/product/88260#/88260. Accessed 4 Dec 2019 2019

