## Supplementary Information for "Fully Automated Peptide Mapping Protocol for Multi-Attribute Method by Liquid Chromatography–Tandem Mass Spectroscopy with a High-Throughput Robotic Liquid Handling System"

### **Design of Experiments**

The design of experiments (DoE) was conducted under RStudio environment with R DOE packages installed. Given the experimental variables, which are microdialysis-associated factors, randomized screening experiments with 32 runs were generated (Table S1) to determine factors that were significant in affecting the dialysis efficiency and hence digestion completeness. For simplicity, all screening DoE were conducted using a NISTmAb followed by manual 3.5-hour tryptic digestion.

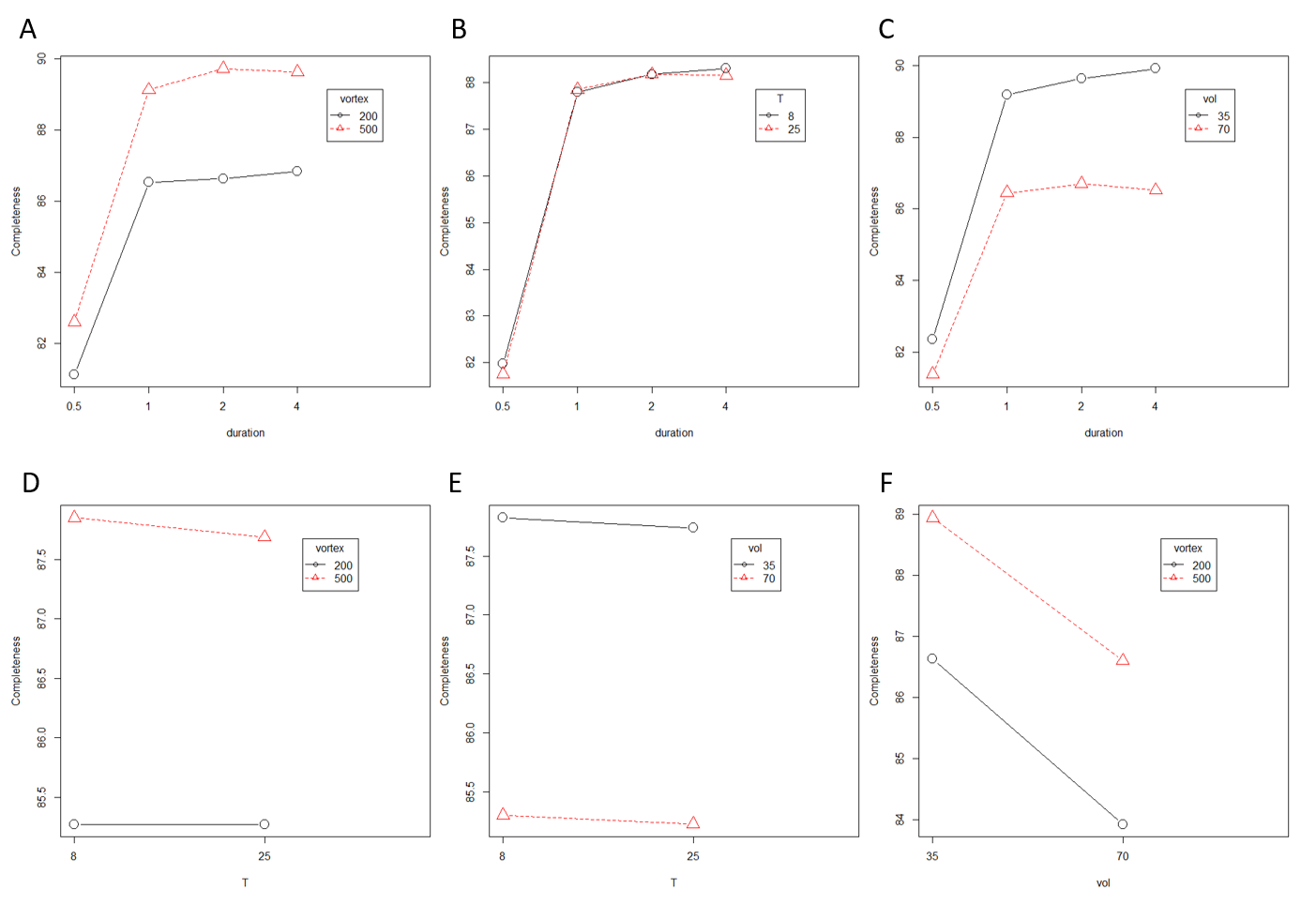

**Figure S1.** Interaction plots by screening DoE to evaluate microdialysis conditions. The effects of four factors—microdialysis duration (0.5, 1, 2, and 4 hours), microdialysis temperature (8 vs 25^o^C), sample volume inside the dialysis cassette (35 vs 70 µL), and microdialysis vortex speed (200 vs 500 rpm)—on digestion completeness were assessed. Digestion completeness increased dramatically with duration from 0.5 hour to 1 hour; however, after 1 hour the dialysis efficiency seemed to reach a plateau (A–C). A vortex speed of 500 rpm during microdialysis yielded noticeably more completeness of digestion than 200 rpm (A, D, F). Similarly, smaller volume inside the dialysis cassette (35 µL) generated a more complete digestion profile (C, E, F). The temperatures during microdialysis did not seem to affect the outcome (B, D, E).

**Table S1.** Randomized screening DoE experimental conditions and digestion completeness results.

| **Run** | **Microdialysis duration (h)** | **Temperature**  **(^o^C)** | **Volume**  **(µL)** | **Vortex speed (rpm)** | **Digestion Completeness (%)** |
| --- | --- | --- | --- | --- | --- |
| 1 | 4 | 8 | 70 | 500 | 88.2 |
| 2 | 2 | 25 | 70 | 200 | 85.1 |
| 3 | 4 | 25 | 35 | 500 | 91.0 |
| 4 | 4 | 25 | 70 | 500 | 88.0 |
| 5 | 4 | 8 | 70 | 200 | 84.8 |
| 6 | 2 | 8 | 70 | 500 | 88.5 |
| 7 | 0.5 | 8 | 70 | 200 | 81.0 |
| 8 | 2 | 8 | 70 | 200 | 84.9 |
| 9 | 1 | 25 | 70 | 200 | 84.7 |
| 10 | 4 | 25 | 70 | 200 | 85.1 |
| 11 | 0.5 | 8 | 70 | 500 | 81.9 |
| 12 | 4 | 25 | 35 | 200 | 88.5 |
| 13 | 2 | 8 | 35 | 500 | 91.2 |
| 14 | 4 | 8 | 35 | 200 | 88.9 |
| 15 | 1 | 25 | 35 | 200 | 88.4 |
| 16 | 0.5 | 8 | 35 | 500 | 83.5 |
| 17 | 1 | 25 | 35 | 500 | 90.3 |
| 18 | 2 | 25 | 35 | 200 | 88.4 |
| 19 | 1 | 8 | 35 | 200 | 88.0 |
| 20 | 0.5 | 25 | 70 | 500 | 81.8 |
| 21 | 2 | 25 | 70 | 500 | 88.3 |
| 22 | 0.5 | 25 | 35 | 500 | 83.2 |
| 23 | 2 | 25 | 35 | 500 | 90.9 |
| 24 | 0.5 | 25 | 35 | 200 | 81.2 |
| 25 | 0.5 | 8 | 35 | 200 | 81.5 |
| 26 | 1 | 25 | 70 | 500 | 88.0 |
| 27 | 4 | 8 | 35 | 500 | 91.3 |
| 28 | 2 | 8 | 35 | 200 | 88.1 |
| 29 | 1 | 8 | 70 | 200 | 85.0 |
| 30 | 1 | 8 | 70 | 500 | 88.1 |
| 31 | 0.5 | 25 | 70 | 200 | 80.8 |
| 32 | 1 | 8 | 35 | 500 | 90.1 |

### **Dialysis cassette position calibration**

One of the major challenges of automating the 96-well microdialysis-based workflow was to navigate the robotic arms precisely to the 1-mm-diameter port of each microdialysis device and execute the sample dispensing and recovery steps. To do this, we leveraged the Hamilton Heater Shaker (Hamilton Company, Reno, NV) and its built-in plate clamp as an active plate nest, fixing the position of the microdialysis plate on the deck relative to the pipetting arm. Fixing the plate position in this way minimized the positional variability of the plate and ensured that the plate position lay well within the 1-mm position tolerance needed for the ports of the dialysis device. From here, iterative tuning of the x, y, and z dimensions of the dialysis plate in the Venus software enables the alignment of the pipetting channel precisely to the dialysis port to maximize the tip seal and ensure proper sample dispensing and recovery.

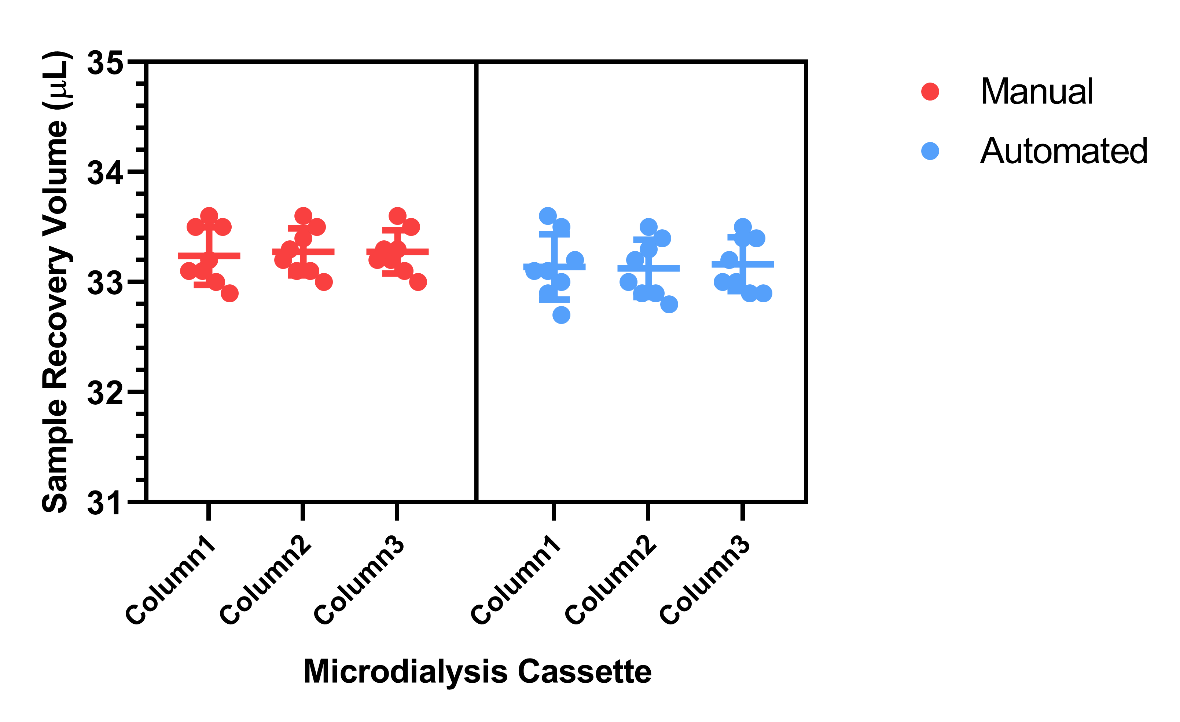

**Figure S2.** Comparison of sample volumes recovered from microdialysis cassettes (35 μL inside each device) in the manual and automated procedures. The nonrecoverable volume inside each device was approximately 2 μL. The total sample volumes recovered were comparable between the manual and the automated procedure.

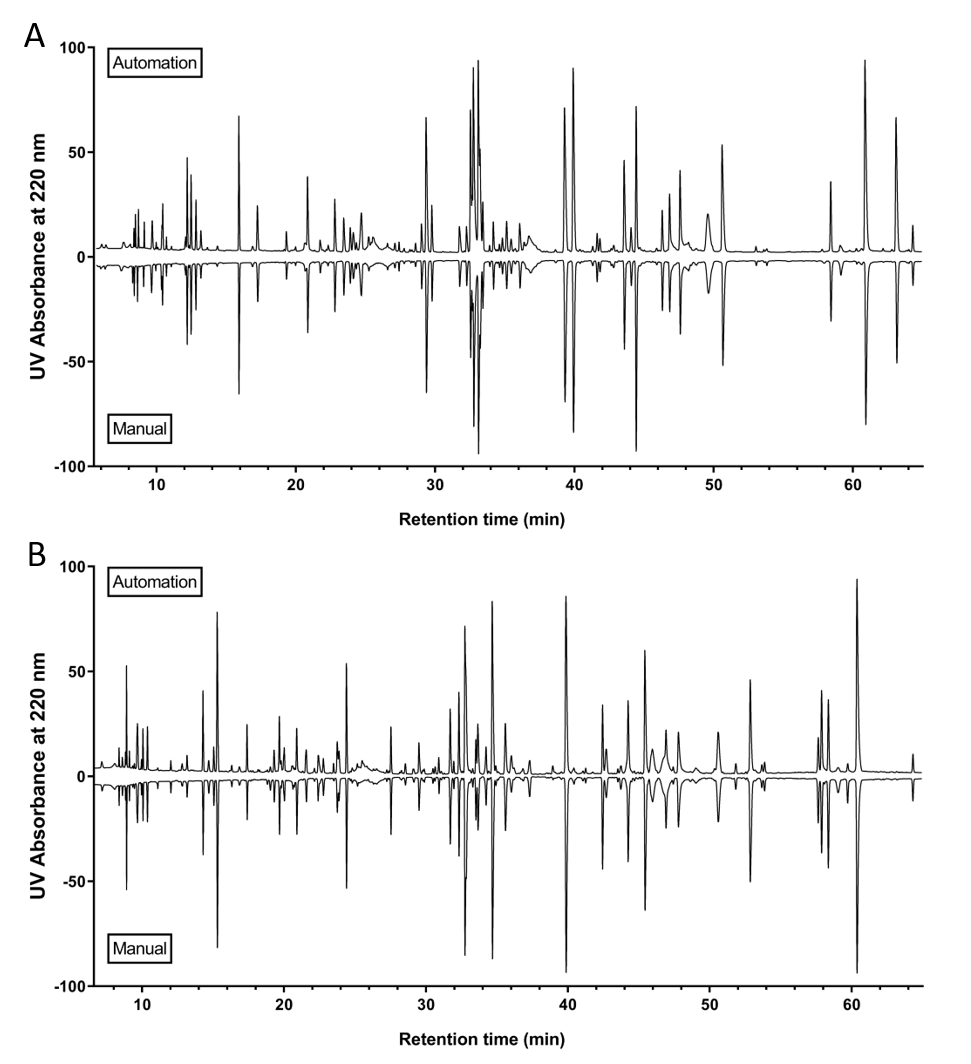

**Figure S3.** Butterfly plot of UV chromatograms comparing the tryptic peptide mapping digestion profiles of an IgG1 (A) and an IgG2 monoclonal antibody. The profiles generated from the automated procedure are displayed on the top and the manual at the bottom. For both mAbs, the digestion was complete with major peptides covering the sequence of 99% of heavy chain, and 100% of light chain. The digestion profiles from automated procedure are highly comparable to those generated from the manual procedure.

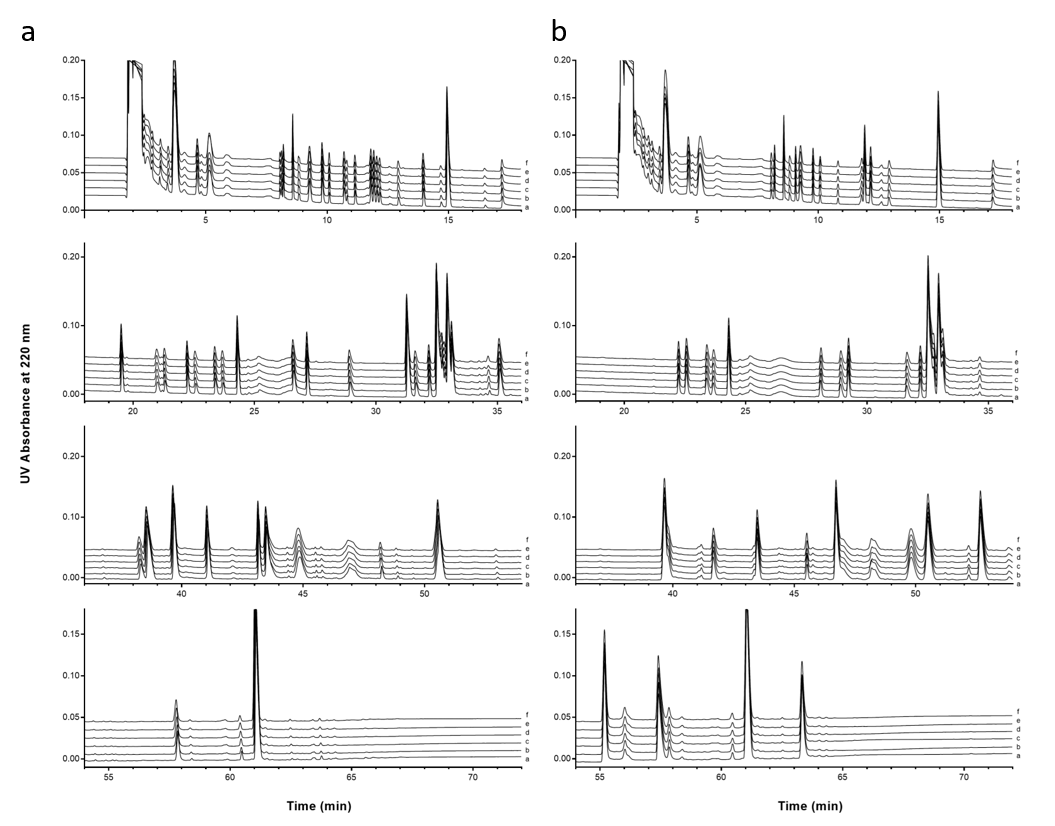

**Figure S4.** UV chromatogram overlay from six replicates of (a) an IgG1 monoclonal antibody and (b) an IgG2 monoclonal antibody.
